# Mapping splice QTLs reveals distinct transcriptional and post-transcriptional regulatory variation of gene expression in pigs

**DOI:** 10.1101/2022.11.20.517281

**Authors:** Fei Zhang, Deborah Velez-Irizarry, Catherine W Ernst, Wen Huang

**Affiliations:** Department of Animal Science, Michigan State University, East Lansing, MI 48824

**Keywords:** splice QTL, expression QTL, regulatory variation of gene expression, pigs

## Abstract

**Background:** Alternative splicing is an important step in gene expression, generating multiple isoforms for the same genes and greatly expanding the diversity of proteomes. Genetic variation in alternative splicing contributes to phenotypic diversity in natural populations. However, the genetic basis of variation in alternative splicing in livestock animals including pigs remains poorly understood.

**Results:** In this study, using a Duroc x Pietrain F2 pig population, we performed genome-wide analysis of alternative splicing estimated from stranded RNA-Seq data in skeletal muscle. We characterized the genetic architecture of alternative splicing and compared its basic features with overall gene expression. We detected a large number of novel alternative splicing events that were not previously annotated. We found heritability of quantitative alternative splicing scores (percent spliced in or PSI) to be lower than that of overall gene expression. In addition, heritabilities showed little correlation between alternative splicing and overall gene expression. Finally, we mapped expression QTLs (eQTLs) and splice QTLs (sQTLs) and found them to be largely non-overlapping.

**Conclusions:** Our results suggest that regulatory variation exists at multiple levels and that their genetic controls are distinct, offering opportunities for genetic improvement.

## Background

Expression of genes is regulated by a multitude of molecular processes which collectively determine the abundance and spatiotemporal distribution of gene expression products of different forms. Both genetic and environmental factors can lead to variation in gene expression regulation, which can ultimately cause variation in organismal phenotypes within and between species. For example, it has been well established that variation in gene expression contributes to the divergence between species [1]. Recent genome-wide mapping efforts have identified numerous expression quantitative trait loci (eQTLs), *i*.*e*. DNA variants that are associated with changes in gene expression, within different organisms, including plants [2], humans [3], insects [4], and livestock animals [5].

Alternative splicing of pre-mRNA is a major step in post-transcriptional regulation of complex eukaryotic gene expression. The most prominent consequence of alternative splicing is the vast expansion of the proteome by forming multiple transcript isoforms, which have the potential to code for different proteins [6]. Alternative splicing is tightly controlled, often in a cell type and developmental stage specific manner, to regulate fundamental biological processes including growth and development. Despite the importance of alternative splicing in gene expression regulation, the genetic basis of alternative splicing variation between and within species and its relationship with phenotypic variation remain less understood than steady state RNA abundances.

Importantly, genetic variation in alternative splicing is abundant and contributes significantly to phenotypic variation in natural populations [7]. For example, aberrant alternative splicing caused by mutations in *cis* regulatory elements or splicing factors underlies many human diseases [8–10]. In livestock animals, QTLs affecting alternative splicing have been mapped in dairy and beef cattle and found to be extensively shared across tissues [11]. Many tissue specific QTLs influencing alternative splicing were also present [11]. In addition, eQTLs and splice QTLs (sQTLs) were found to be associated with meat quality traits in a population of Angus-Brahman crossbred cattle [12].

In a previous study, we mapped hundreds of eQTLs for RNA abundance in the *longissimus dorsi* muscle of pigs from a Duroc x Pietrain F2 population, providing potential links to facilitate the functional interpretation and dissection of phenotypic QTLs [5]. In the present study, we extended this work to map sQTLs, taking advantage of the paired-end and strand-specific RNA-Seq data. We compared the heritability of overall gene expression and alternative splicing which showed little correlation. Furthermore, the comparison of mapped eQTLs and sQTLs showed little overlap and different location patterns, suggesting distinct transcriptional and post-transcriptional regulatory variation of gene expression.

## Results

### Identification of alternative splicing events in pigs using RNA-Seq

To understand the genetic basis of alternative splicing regulation in skeletal muscle in pigs, we analyzed the RNA-Seq dataset from a previous study [5] to obtain both overall gene expression and alternative splicing quantifications in *longissimus dorsi* muscle. We focused on 143 of the 168 animals for which strand-specific RNA-Seq libraries were sequenced. On average, 45.8 million 125 bp paired-end reads were sequenced per sample, 37.5 million of which (Additional File 1) were uniquely mapped using HISAT2 [13]. Importantly, of the uniquely mapped reads, 21.5 million covered junction sites, enabling us to classify and quantify alternative splicing (Additional File 1).

We assembled transcript models from each sample using StringTie [14] and merged and filtered them to produce a combined annotation. We identified alternative splicing events from merged transcript annotations across all RNA-Seq samples using rMATs [15], which classified them into five classes, including skipped exon (SE), retained intron (RI), alternative 5′ splice site (A5SS), alternative 3′ splice site (A3SS), and mutually exclusive exons (MXE). We considered an alternative splicing event supported by experimental evidence in our muscle RNA-Seq data if both splice isoforms were supported by at least two junction reads in at least two samples. There were 88,670, 4,980, 3,044, 3,591, and 23,187 SE, RI, A5SS, A3SS, and MXE events respectively, among which, 7,323, 2,285, 1,720, 1,963, and 2,081 were novel (Figure 1). Despite that only muscle RNA was sequenced, the large number of unannotated alternative splicing events testified to the power of population-scale RNA-Seq and the incompleteness of the current gene annotation in pigs.

**Figure 1.**
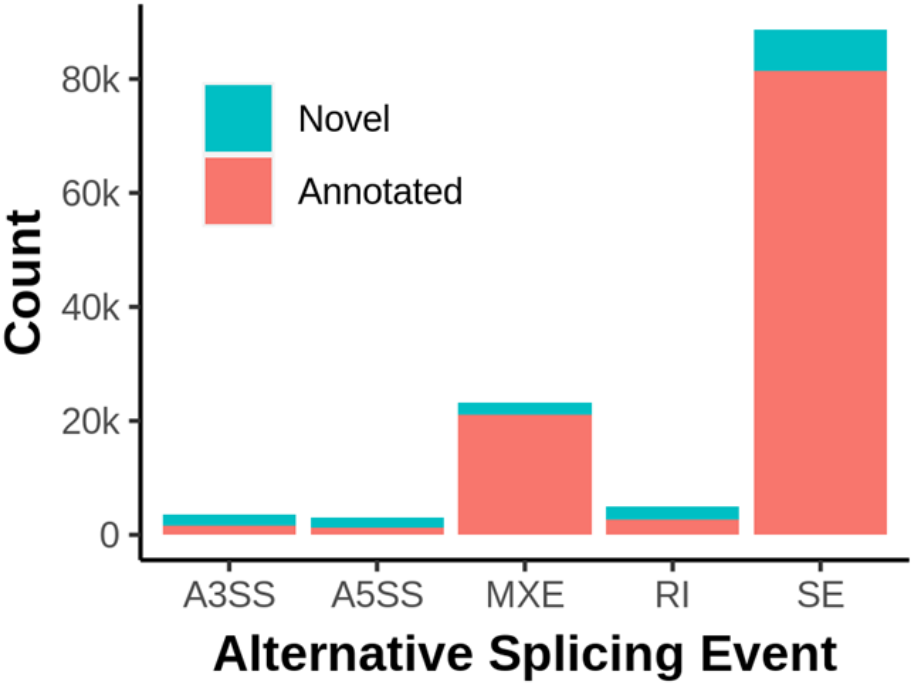
Identification of annotated and novel alternative splicing events in pig muscle by RNA-Seq. Stacked bar chart showing the counts of annotated and novel alternative splicing events supported by experimental evidence. We consider an event experimentally supported if both isoforms are supported by at least two reads in at least two samples. A3SS = alternative 3’ splice site; A5SS = alternative 5’ splice site; MXE = mutually exclusive exons; RI = retained intron; SE = skipped exon.

### Quantitative genetics of alternative splicing and steady state RNA abundance

To understand the genetic basis of alternative splicing and steady state RNA abundance, we first estimated the genomic heritability of these two types of traits. Genomic heritability quantifies the proportion of variation in alternative splicing or overall gene expression that can be attributed to additive genetic variance. We defined the percent spliced-in (PSI) score as the proportion of junction reads supporting one of the splice isoforms versus the sum of both isoforms to quantitatively represent alternative splicing.

We applied several non-specific filters to PSI and TPM (transcripts per million) representing overall gene expression, including filters on overall magnitude of variances, coefficient of variation and skewness of distribution. The remaining genes or alternative splicing events were Normal quantile transformed and fitted in a linear mixed model with genomic relationship matrix calculated from 50K SNP chip genotypes of the animals. Significance of the deviation of additive genetic variance from zero was determined using a likelihood ratio test comparing the full mixed model and a reduced model without the additive genetic variance component. Among a total of 11,368 genes and 22,018 alternative splicing events in 6,004 genes, 2,200 genes and 405 alternative splicing events in 190 genes had significant heritability at a false discovery rate (FDR) = 0.01. A total of 65 genes had both significant heritable overall gene expression and alternative splicing.

The estimated heritability was in general larger for overall gene expression than for all five types of alternative splicing (Figure 2). Importantly, the heritability of these two aspects of gene expression showed little correlation (Figure 3). Bivariate quantitative genetic analysis suggested that the genetic correlation was generally small. Of the 20,625 gene expression and alternative splicing pairs, only 585 had a significant (FDR = 0.01) genetic correlation.

**Figure 2.**
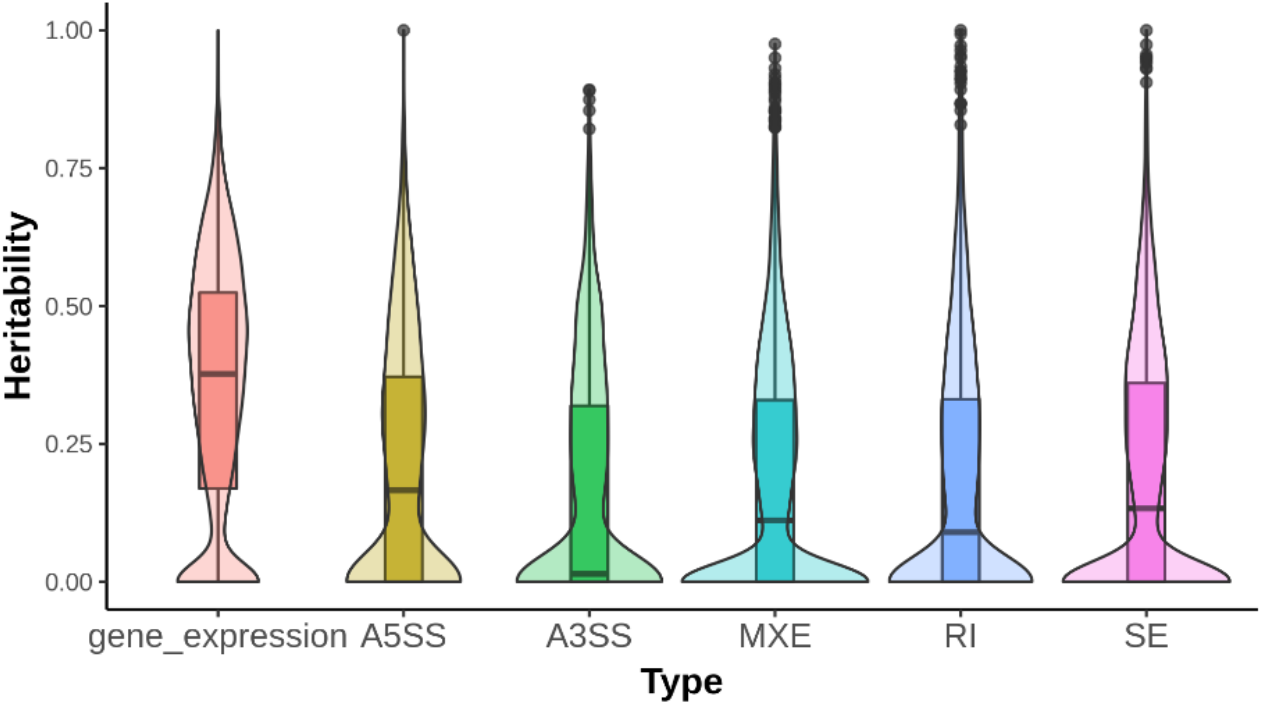
Distribution of genomic heritabilities for gene expression and alternative splicing traits. Violin plots overlaid on boxplots showing the distribution of genomic heritability for overall gene expression and five classes of alternative splicing. A3SS = alternative 3’ splice site; A5SS = alternative 5’ splice site; MXE = mutually exclusive exons; RI = retained intron; SE = skipped exon.

**Figure 3.**
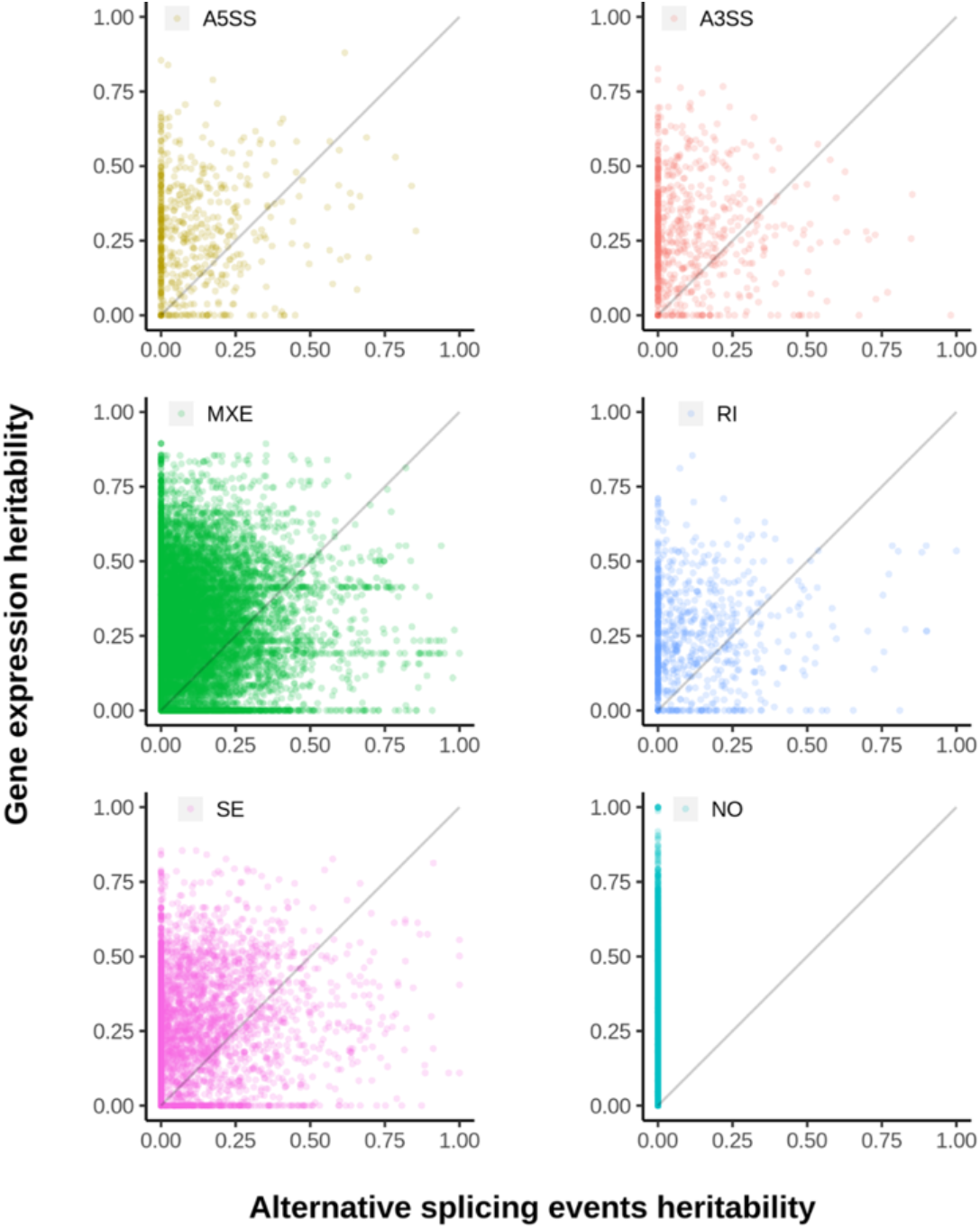
Comparison of heritability for alternative splicing and overall gene expression for the same genes. Scatter plots showing comparison between heritability for five classes of alternative splicing and overall gene expression. The diagonal line indicates perfect agreement. A5SS = alternative 5’ splice site; A3SS = alternative 3’ splice site; MXE = mutually exclusive exons; RI = retained intron; SE = skipped exon; NO = no alternative splicing events detected. Multiple events may exist for a gene and are represented by different points on the plots.

These results suggested that the heritable genetic control of transcriptional and post-transcriptional regulation of gene expression may be distinct, even though the two molecular processes are highly coupled.

### Mapping splice QTLs (sQTLs) and expression QTLs (eQTLs)

To understand the distinct genetic control of transcription and splicing, we mapped splice QTLs (sQTLs) and expression QTLs (eQTLs) for all genes using genome-wide association analysis. At an empirical FDR = 0.01, we identified a total of 35,687 sQTLs for 796 alternative splicing events in 475 genes and 43,432 eQTLs for 1,098 genes (Figure 4a,b). The large number of sQTLs and eQTLs per alternative splicing or gene expression trait was the result of the high level of linkage among genetic markers in the F2 population. We therefore performed a forward model selection approach to eliminate associations due to linkage and classified sQTLs and eQTLs based on the markers retained in the models. Both sQTLs and eQTLs exhibited an enrichment of *cis* (defined as 1Mb within the boundaries of genes) QTLs as compared to *trans*. Despite that *trans* QTLs have a much larger space to exist, of the 1,338 genes with at least one eQTL and 961 alternative splicing events with at least one sQTL, 552 (41%) and 467 (48%) had *cis* eQTLs and sQTLs, respectively (Table 1). These results, as many studies have shown for eQTLs previously, indicated that proximal regulation is a primary mode of regulation for both transcription and alternative splicing.

**Table 1.** Counts of eQTLs and sQTLs classified into *cis* and *trans*

|  | **Total** | ***cis*** | ***trans*** | **both** |
| --- | --- | --- | --- | --- |
| Genes | 1,338 | 552 | 865 | 79 |
| Alternative splicing events | 961 | 467 | 549 | 55 |

**Figure 4.**
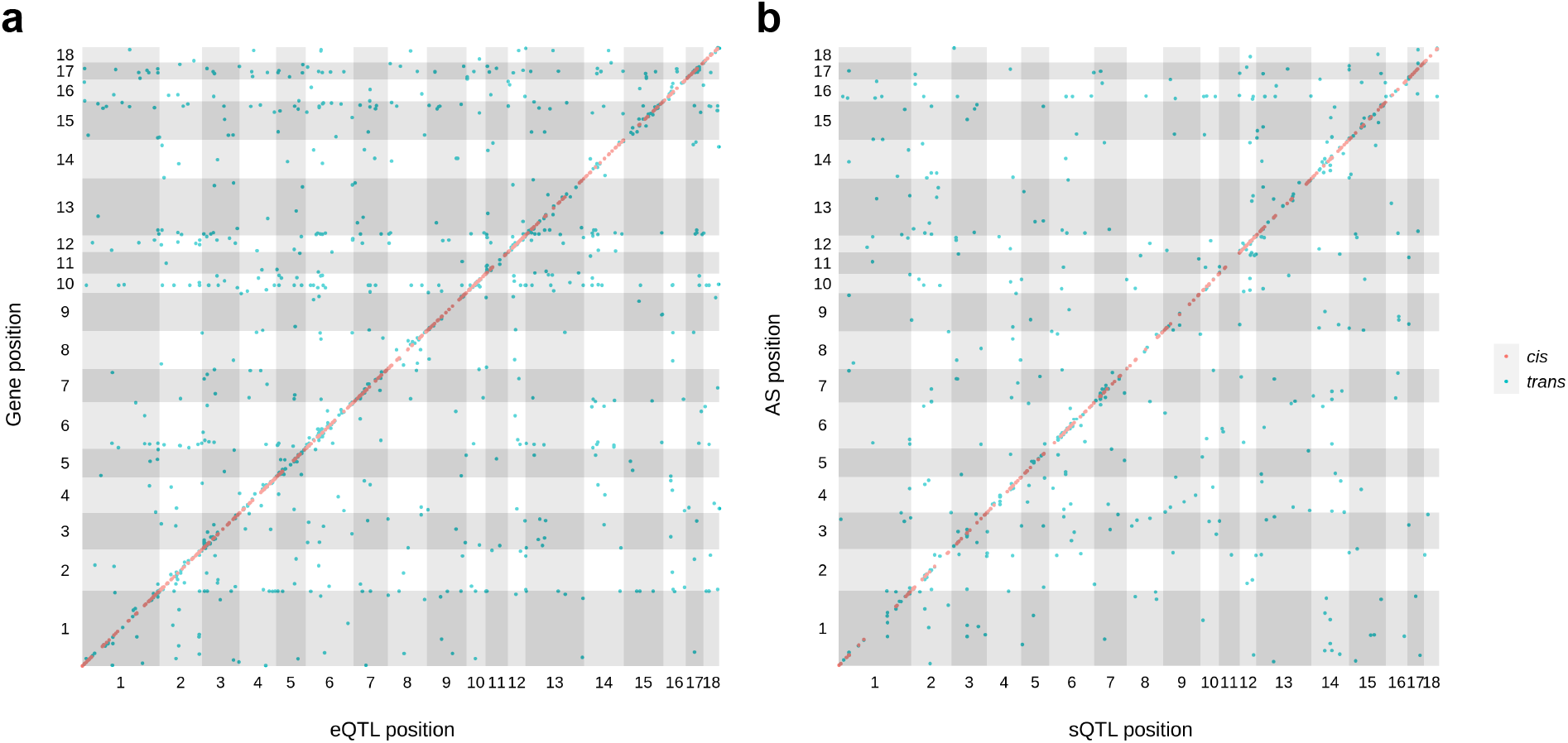
Locations of eQTLs and sQTLs relative to genes. eQTL (a) and sQTL (b) location plots showing the relative positions of eQTLs and sQTLs versus genes and alternative splicing events. Red and green dots indicate cis and trans QTLs respectively. Chromosomes are shown using alternative grey and white colors.

Among the 1,338 genes with eQTLs and 586 genes with sQTLs, 105 were in common. Of the 105 genes with mapped eQTLs and sQTLs, only 16 contained SNPs that were associated with both overall gene expression and alternative splicing. These results, together with the lack of correlation between SNP based heritability estimates for overall gene expression and alternative splicing and low genetic correlation, suggested that the genetic control of transcription and alternative splicing were likely distinct.

## Discussion

In this study, we carried out an analysis to estimate heritability of alternative splicing and map sQTLs in skeletal muscle of 143 pigs in a Duroc x Pietrain F2 population [5]. The samples were deeply sequenced with paired-end reads, enabling us to estimate both alternative splicing and steady state RNA abundances. While many studies have mapped eQTLs for overall gene expression in livestock animals, genetic characterization of alternative splicing remains understudied.

With a single tissue type, we found a significant number of novel alternative splicing events supported by multiple reads in multiple samples (Figure 1). Skeletal muscle is one of the most intensively studied tissues in pigs and yet we were able to substantially improve existing annotations, underscoring the incompleteness of the present annotation, which is expected to further improve as large-scale annotation projects are completed [16].

By comparing genomic heritability of quantitative gene expression and alternative splicing, we found heritability of alternative splicing was generally lower than that of overall expression (Figure 2). It’s important to point out that the magnitude of heritability depends on the precision of measurements. An explanation for the lower heritability for alternative splicing may be that alternative splicing utilizes only reads spanning junctions, whereas overall gene expression takes advantage of all exonic reads.

We found little correlation between the heritabilities of overall gene expression and alternative splicing (Figure 3). This suggested that genetic controls of transcription and alternative splicing were likely distinct. In addition, we estimated generally low genetic correlation between overall expression and alternative splicing with high standard errors. Out of 20,625 gene expression and alternative splicing pairs, only 585 had a significant (FDR = 0.01) genetic correlation. Both eQTLs and sQTLs were enriched for SNPs proximal to the genes they regulated, where *cis* regulatory elements were concentrated. A substantial fraction of genes contained both mapped eQTLs and sQTLs, yet only a small fraction shared the same genetic markers as eQTLs and sQTLs. The resolution of the F2 design and the density of the SNP chip did not allow us to perform a comprehensive evaluation of overlap. Taken together, these results suggested that transcription and alternative splicing were distinctly regulated genetically.

Ultimately, regulatory variation in transcription and splicing lead to phenotypic variation. Our study suggests that when considering regulatory variation as an intermediate step that connects DNA variation and organismal phenotypic variation, alternative splicing should not be ignored. Indeed, studies in humans suggest that alternative splicing may be a major contributor to phenotypic variation [7]. As large-scale annotation projects improve annotation of the pig genome, estimation of alternative splicing will become easier and more accurate, which will ultimately improve our ability to understand and utilize its contribution to phenotypic variation.

## Conclusions

Using a stranded RNA-Seq dataset in a F2 population of pigs, we mapped eQTL and sQTL in the skeletal muscle. We found overall gene expression and alternative splicing to have low genetic correlation, and heritabilities of these two classes of traits to have low correlation. In addition, although both classes of traits had QTLs concentrated near the genes, we found very little overlap between sQTLs and eQTLs. These results suggested that the genetic regulation of transcription and alternative splicing were likely distinct that both need to be considered when mapping regulatory variation that connects DNA variation and phenotypic variation.

## Materials and Methods

### RNA-Seq data mapping and alternative splicing event calling

All RNA-Seq data were obtained from a previously published study with accession number PRJNA403969 [5]. To reduce the influence of batch effect, we focused on 144 samples which were prepared uniformly to produced stranded sequencing libraires and sequenced by the Illumina HiSeq 2500 platform with 125 bp paired end reads. Raw sequence reads were aligned to the *Sus scrofa* reference genome (11.1) and transcriptome (Ensembl Release 97) using HISAT2 2.1.0 [13]. There were on average 45.8 million of reads per sample, with 90.9% overall mapping and 81.9% unique mapping rates. One of the samples (1833) had an uncharacteristically low (49.64%) unique mapping rate and was removed from further analysis, resulting in 143 samples. Mapped reads that covered junctions comprised 46.9% of all reads on average, providing a strong basis to estimate alternative splicing accurately.

StringTie 2.04 [14] was used to assemble transcripts based on HISAT2 alignments, producing a gene annotation per sample. The StringTie parameters were set to allow a minimum transcript length of 200 bp, minimum transcript coverage of 2.5 and minimum isoform fraction of 0.1. Annotations from all samples were merged to a set of 78,351 transcripts, among which 29,193 (37.26%) were novel. Alternative splicing events were identified by rMATs 4.1.0 [15] using the combined gene annotations and novel alternative splicing was called if they did not exist in the reference annotation. We only considered novel events where both isoforms were supported by at least two reads in two samples.

### Heritability estimation of alternative splicing and gene expression

To quantify alternative splicing, percent splice-in (PSI) scores [15] were calculated for each gene using the following formula:

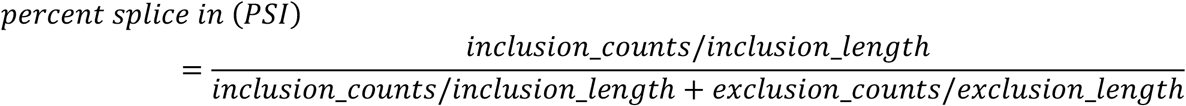

When estimating heritability, we only considered alternative splicing events with at least 5 total reads supporting both isoforms in at least 50% of the samples. Only alternative splicing events with a mean absolute deviation greater than 1, log2 variance greater than 2, skewness < 1.1 were considered variable and retained. In addition, to eliminate the influence of sex specific alternative splicing, we only retained events where samples from either sex contributed at least 20% of the estimable samples. PSI was normal quantile transformed using the R package bestNormalize [17]. The transformed PSI was used to estimate genomic heritability (*h*^*2*^) using the R package rrBLUP [18] with sex of the animal as a fixed effect. The R package fdrtool was then used to calculate the FDR value based on *P* values obtained from likelihood ratio tests.

Overall gene expression was estimated using transcript per million (TPM) output by StringTie. TPM of each gene was also subjected to a normal quantile transformation. The transformed TPM was then used to estimate the genomic heritability (h^2^) of gene expression by rrBLUP package. Genetic correlation was computed using the GCTA package [19]. The R package fdrtool was then used to calculate the FDR values based on the Benjamini-Hochberg approach.

### Splicing QTL and expression QTL mapping

eQTL and sQTL mapping was performed using the fastGWA function of GCTA [19] with sex of the animal fitted as a covariate. To estimate empirical FDR, we permuted the data set (animal ID) 200 times while preserving correlation among genes and performed eQTL and sQTL mapping in the permuted datasets. Empirical FDR for each was estimated as the observed number of significant QTLs at a given threshold divided by the average number of significant QTLs among the 200 permutations. eQTLs or sQTLs located within 1Mb of gene boundaries were consider *cis*, and otherwise *trans*.

## Authors’ contributions

F.Z., D.V. C.W.E. and W.H. conceived and designed the study; F.Z. analyzed the data; D.V. and C.W.E. contributed analytical tools; F.Z. and W.H. wrote the manuscript with input from all authors. All authors read and approved the submitted version.

## Code availability

Codes are available at: https://github.com/zhangfei1947/MSUPRP.

## Ethics approval and consent to participate

Not applicable.

## Consent for publication

Not applicable.

## Reference

1. King M, Wilson A. Evolution at two levels in humans and chimpanzees. Science (1979). 1975;188:107–16.

2. Zhang H, Mao R, Wang Y, Zhang L, Wang C, Lv S, et al. Transcriptome-wide alternative splicing modulation during plant-pathogen interactions in wheat. Plant Sci. 2019;288.

3. Aguet F, Brown AA, Castel SE, Davis JR, He Y, Jo B, et al. Genetic effects on gene expression across human tissues. Nature. 2017;550:204–13.

4. Huang W, Carbone MA, Magwire MM, Peiffer JA, Lyman RF, Stone EA, et al. Genetic basis of transcriptome diversity in Drosophila melanogaster. Proceedings of the National Academy of Sciences. 2015;112:E6010–9.

5. Velez-Irizarry D, Casiro S, Daza KR, Bates RO, Raney NE, Steibel JP, et al. Genetic control of longissimus dorsi muscle gene expression variation and joint analysis with phenotypic quantitative trait loci in pigs. BMC Genomics. 2019;20.

6. Nilsen TW, Graveley BR. Expansion of the eukaryotic proteome by alternative splicing. Nature. 2010;463:457–63.

7. Li YI, van de Geijn B, Raj A, Knowles DA, Petti AA, Golan D, et al. RNA splicing is a primary link between genetic variation and disease. Science (1979). 2016;352:600–4.

8. Waldron D. Human genetics: Splicing: linking genetic variation and disease. Nat Rev Genet. 2016;17:317.

9. Sterne-Weiler T, Sanford JR. Exon identity crisis: disease-causing mutations that disrupt the splicing code. Genome Biol. 2014;15.

10. Garcia-Blanco MA, Baraniak AP, Lasda EL. Alternative splicing in disease and therapy. Nat Biotechnol. 2004;22:535–46.

11. Xiang R, Hayes BJ, vander Jagt CJ, MacLeod IM, Khansefid M, Bowman PJ, et al. Genome variants associated with RNA splicing variations in bovine are extensively shared between tissues. BMC Genomics. 2018;19.

12. Leal-Gutiérrez JD, Elzo MA, Mateescu RG. Identification of eQTLs and sQTLs associated with meat quality in beef. BMC Genomics. 2020;21.

13. Kim D, Paggi JM, Park C, Bennett C, Salzberg SL. Graph-based genome alignment and genotyping with HISAT2 and HISAT-genotype. Nat Biotechnol. 2019;37:907–15.

14. Pertea M, Pertea GM, Antonescu CM, Chang T-C, Mendell JT, Salzberg SL. StringTie enables improved reconstruction of a transcriptome from RNA-seq reads. Nat Biotechnol. 2015;33:290–5.

15. Shen S, Park JW, Lu Z, Lin L, Henry MD, Wu YN, et al. rMATS: Robust and flexible detection of differential alternative splicing from replicate RNA-Seq data. Proceedings of the National Academy of Sciences. 2014;111:E5593–601.

16. Andersson L, Archibald AL, Bottema CD, Brauning R, Burgess SC, Burt DW, et al. Coordinated international action to accelerate genome-to-phenome with FAANG, the Functional Annotation of Animal Genomes project. Genome Biol. 2015;16:57.

17. Peterson RA, Cavanaugh JE. Ordered quantile normalization: a semiparametric transformation built for the cross-validation era. J Appl Stat. 2019;47:2312–27.

18. Endelman JB. Ridge Regression and Other Kernels for Genomic Selection with R Package rrBLUP. Plant Genome. 2011;4:250–5.

19. Yang J, Lee SH, Goddard ME, Visscher PM. GCTA: A tool for genome-wide complex trait analysis. Am J Hum Genet. 2011;88:76–82.

